# Explore synergistic and competitive miRNA regulation mechanisms in the miRNA-mRNA regulatory network from the information decomposition perspective

**DOI:** 10.1101/2021.12.20.473520

**Authors:** Chu Pan, Jing Jiang, Limei Jing, Wangqing Chen, Yi Yang, Ying Liu, Jiawei Luo, Xiangxiang Zeng

## Abstract

Since multiple microRNAs can target 3’ untranslated regions of the same mRNA transcript, it is likely that these endogenous microRNAs may form synergistic alliances, or compete for the same mRNA harbouring overlapping binding site matches. Synergistic and competitive microRNA regulation is an intriguing yet poorly elucidated mechanism. We here introduce a computational method based on the multivariate information measurement to quantify such implicit interaction effects between microRNAs. Our informatics method of integrating sequence and expression data is designed to establish the functional correlation between microRNAs. To demonstrate our method, we exploited TargetScan and The Cancer Genome Atlas data. As a result, we indeed observed that the microRNA pair with neighbouring binding site(s) on the mRNA is likely to trigger synergistic events, while the microRNA pair with overlapping binding site(s) on the mRNA is likely to cause competitive events, provided that the pair of microRNAs has a high functional similarity and the corresponding triplet presents a positive/negative ‘synergy-redundancy’ score.

## INTRODUCTION

MicroRNA (miRNA) is an endogenous noncoding RNA that regulates gene expression post-transcriptionally by pairing to mRNAs at the 3’ untranslated regions (UTR)(1). The binding event primarily occurs at the 2-8 nucleotide positions from the 5’ end of the miRNA, which is termed as the seed region; the binding is termed as seed matching(2,3). The composition of seed sequence is the major determinant of miRNA targeting patterns(4). Since the miRNA binding sites on the 3’ UTR of the same RNA transcript may be neighbouring or overlapping, there are likely to cause synergistic or competitive interaction effects between miRNAs (Figure 1A). Indeed, the coordinating actions of miRNAs are rampant in different cell phenotypes especially in cancer pathologies(5,6), a classic example being the cooperative effects of miR-200bc/429 cluster in breast cancer progression(7). Alternatively, antagonism between miRNAs was observed in the transfected cells in case of overlapping binding sites or limited Argonaute(8). These evidences suggest that the interaction effects between miRNAs indeed exist and interfere with the expression level of genes inside cells, which in turn results in profound impact on the occurrence of complex diseases including tumorigenesis. While the internal mechanism of such widespread interaction effects between miRNAs has not been systematically interpreted. Yet both synergistic and competitive relationship have been experimentally supported.

**Fig. 1.**
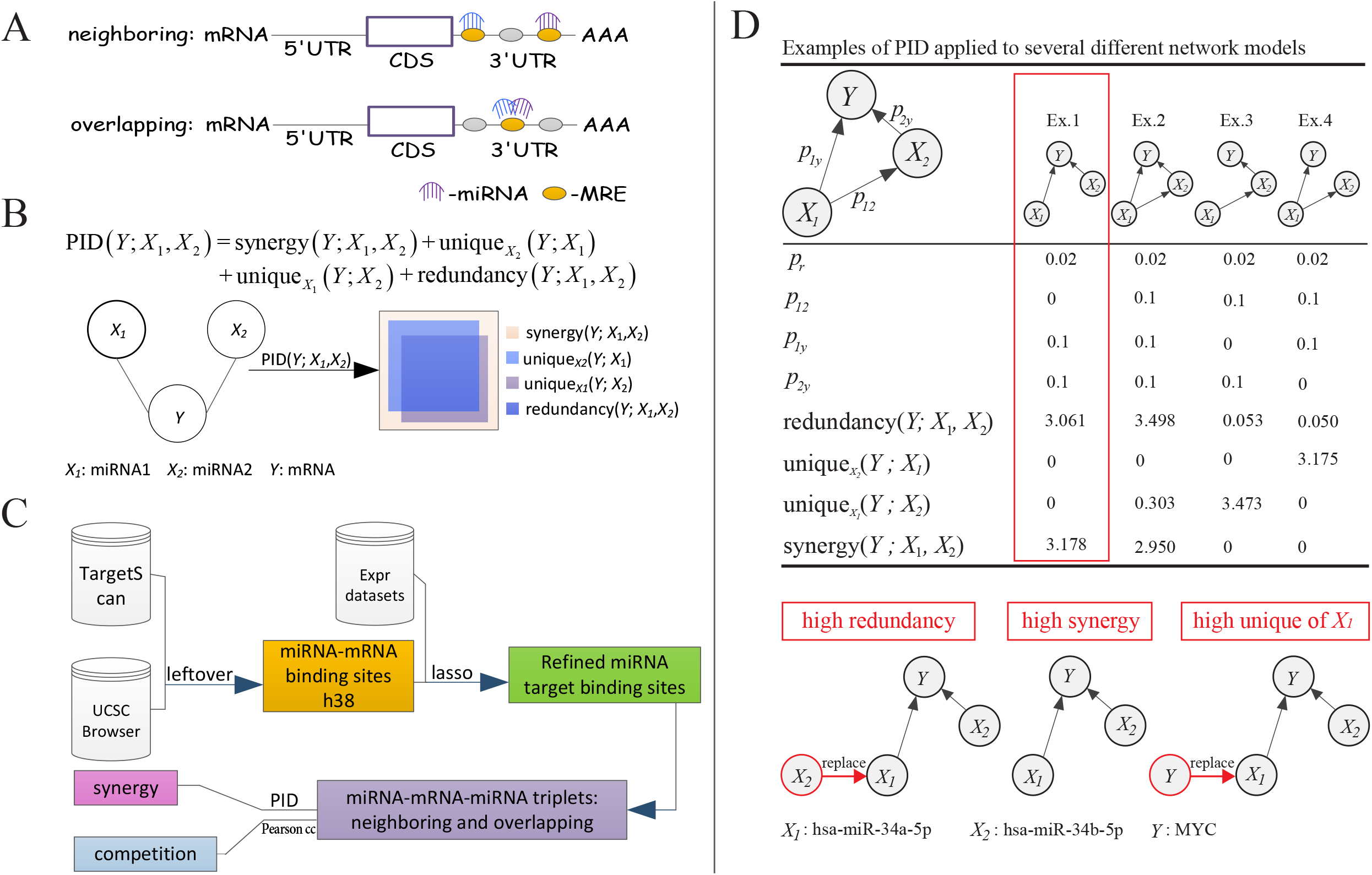
sycomore model overview. **A** Two different scenarios where two miRNAs target the same mRNA transcript. In scenario one (neighbouring), both miRNAs target the same mRNA transcript with neighbouring binding sites. In Scenario two (overlapping), both miRNAs target the same mRNA transcript with overlapping binding sites. **B** Our three-variable network structure. **C** The workflow of sycomore. **D** Information distribution of different three-variable network structures by PID. Figure referred to the work by Timme *et al*. (24). The general structure of the network model, the original probability values, the four variants and the corresponding results were demonstrated in upper of Figure **1D**. The general structure of our network model along with the three alternate combinations of source and target variables were presented in the lower of Figure **1D**.

Perhaps the most extensive computational studies about miRNAs have focused on the prediction of their targets(2,9-15). Among these computational methods, a few studies have considered the potential miRNA interactions into the design of prediction methods. For instance, Krek *et al*. inputted multiple highly co-expressed miRNA sequences into a hidden Markov-based probability model at one time to predict the cumulative probability of binding to the same target. The results showed that combinatorial miRNA target predictions reduced the false positive rates of predictions, and also provided evidence for coordinate miRNA control in mammals(9). On the other hand, Li *et al*. regarded mRNA and miRNA competition as a role switch of the expression seed matching, and thus proposed a probabilistic model to identify miRNA targets. As a target predictor, the proposed model identified high confidence/validated targets(11). In another example, Ding *et al*. simultaneously incorporated the neighbouring and overlapping miRNA binding into a statistical model for miRNA target site prediction. The results showed that using this strategy the proposed model achieved high recall and precision(14). In addition to these miRNA target predictions, Gam *et al*. recently developed a mixed antagonistic/synergistic miRNA repression model to improve the prediction accuracy of multi-input miRNA sensor activity(16). The methods listed above involve miRNA interactions, such as miRNA co-expression or/and miRNA combinatorial binding, in the computational model design. But there still lacks general attempts to address the details of the mechanism behind the synergistic and competitive interaction effects. We thus hypothesize an intuitive and efficient deterministic method to explore such implicit relationship between miRNAs.

In this article, we describe a novel computational method called sycomore to explore the synergistic and competitive interaction effects between two miRNAs when targeting the same mRNA. We integrate sequence and expression data into the method, expecting to fully understand the underlying mechanism of such interaction effects. The main function of sycomore adapts from partial information decomposition (PID), which is a multivariate information measurement that decomposes the Shannon information into a multivariate system in terms of the redundancy between the synergies of source subsets(17). Its role here is to quantify the information that the two joint miRNAs provide about the common target mRNA, and to distinguish between information distribution due to responses of individual miRNAs versus a combination of two miRNAs (Figure 1B). We first refer to the TargetScan data to determine the neighbouring/overlapping of miRNA target site locations. We then test on four different types of cancer data from The Cancer Genome Atlas (TCGA) to detect functional correlation between miRNAs.

## MATERIALS AND METHODS

### MiRNA target site information

The predicted miRNA target sites along with UTR genome coordinates were downloaded from TargetScan Human 7.2(13). The target sites were first mapped into UTR genome coordinates due to the target sites file only provided relative UTR start and end, and then converted the genome coordinates from hg19 to hg38 through the LiftOver tool. The mature miRNA annotation information was downloaded from miRBase database(18). The mRNA annotation information was based on GENCODE human release 34(19). Since most of target mRNA may have multiple alternative mRNA transcripts, we selected the transcript with the longest 3’ UTR as a reference.

### TCGA data collection and pre-processing

We chose four cancer types from the TCGA data portal by considering the proportion of tumour to normal samples and the total sample size. For each cancer type, we downloaded the processed (level 3) RNA-Seq data corresponding to the expression levels of the miRNA and the target mRNA. For the miRNA-Seq data, we chose the mature isoform expression data (i.e., files with extension isoform.quantification.txt) by referring to Li *et al*. (20). We filtered out cross-mapped regions and summed over the RPM (reads per million miRNAs mapped) values and ‘read count’ values for each mature miRNA sequence. For the mRNA-Seq data, we used RPKM (read per kilobase per million mapped reads) values and ‘read count’ values for each mRNA sequence. Both miRNA and mRNA with missing value exceeding 30% of the sample size were filtered out, and the missing value of the remaining miRNAs/mRNAs were all filled with zeros. The remaining expression values were further log2-transformed within each cancer type. In addition, miRNA and mRNA were filtered by differential expression analysis on ‘read count’ value using an R/Bioconductor package DESeq2(21). The miRNAs and mRNAs with | log_2_ FC |≥1 and adj.*P*-value ≤ 0.05 were considered as significantly differentially expressed miRNAs/mRNAs.

### Extraction of cancer-specific miRNA targets via the Lasso regression

We adopted the Lasso regression method, by referring to Li *et al*.(22), to refine the miRNA targets for each cancer via integrating sequence and expression data. The use of Lasso is due to its ability to predict multi-miRNA repression on one mRNA simultaneously, and to cater to cancer-specific miRNA targets. Let us consider a sample consisting of *N* cases, each of which consisting of *M* covariates, that is, the corresponding covariate matrix *X* and the outcome *y*. Its Lagrangian form is as follows:

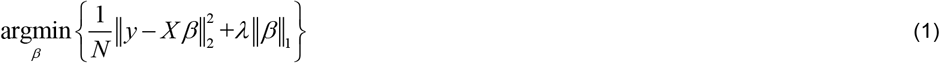

the *y, X* and *β* are the mRNA expression, the miRNA expression and the fitted linear coefficients, respectively, and *λ* is a parameter that controls the strength of L1 penalty. We implemented Lasso using R package glmnet with *α* =1 and extract coefficients at a single value of lambda (i.e., *λ*) by setting *S* = 0.05 in the function coef(23). The fittings in the Lasso regression method are the miRNA expression multiplied by the target sit matrix. Thus, the expression of a miRNA has no effect on mRNA expression if the corresponding target site information matrix entry is zero. Targets with negative correlation are considered as putative functional interactions as the miRNAs are believed to inhibit target expression, and the corresponding miRNA target sites from targetScan are remained.

### Evaluation of information scores for each triplet by partial information decomposition

PID is a recently developed multivariate information measurement introduced by Williams and Beer(17). It aims to investigate the dependence between two or even multiple variables with considerable advantages over the other information measures such as interaction information, as it provides multiple interpretable output items(24). PID can be used to measure the interaction effects between two distinct groups of variables, i.e., a group of source variables and a single variable. In this paper, we are only concerned with the case of three variables. Let *X*_1_, *X*_2_ and *Y* be the first source variable (i.e., miRNA1), the second variable (i.e., miRNA2) and target variable (i.e., mRNA), respectively, and define *R* as the source variable set (Figure 1B). PID decomposes the mutual information provided by the source variables set about a given target variable into four items: synergistic information, two unique information and redundant information. The formula of PID is as follows:

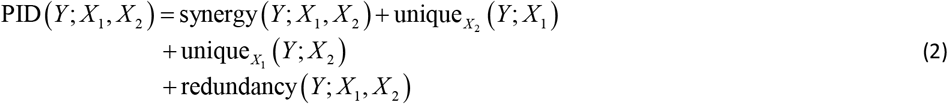

where synergy(*Y*;*X*_1_,*X*_2_) is the partial information about *Y* provided by both source variables in the *R* together, while redundancy(*Y*;*X*_1_,*X*_2_) is the partial information about *Y* provided by either variable in the *R* alone. The unique_x2_(*Y,X*_1_) is the unique information between the source variable *X*_1_ and the target variable *Y* when the other source variable is *X*_*2*_. Similarly, the unique_x1_(*Y,X*_2_) is the unique information between the source variable *X*_2_ and the target variable *Y* when the other source variable is *X*_1_. The structure of PID means that synergistic and redundant interactions are not mutually exclusive, as was the case for the traditional interpretation of the interaction information. Please refer to Willams and Beer for specific calculation details(17).

### Details of the sycomore

Our informatics method of integrating sequence and expression data is to quantify the synergistic and competitive interaction effects between miRNAs with two main stages. Figure 1C depicts the flow chart of our method. Before performing the two stages, we first refined candidate miRNA binding sites on the 3’ UTRs of mRNA transcripts, via a series of data pre-processing including mapping the miRNA target sites to the physical location of the 3’ UTR, transforming the genome coordinates from hg19 to hg38 version, and using Lasso regression model to select cancer-specific miRNA targets. At stage 1, we partition miRNA pairs into two groups of neighbouring and overlapping based on the refined physical binding site locations. These miRNA pairs that have adjacent target sites on the same mRNA transcript are classified into the neighbouring group, and these miRNA pairs that have at least 1 nucleotide target site overlapping on the same mRNA transcript are assigned to the overlapping group. At stage 2, we treat each miRNA pair and its common target mRNA as a miRNA-mRNA-miRNA triplet. We apply PID to each triplet to evaluate its information score list at the expression level and employ Pearson correlation coefficient (Pearson cc) to evaluate the functional similarity between the corresponding miRNA pair as well, which we interpret these as evidence of the hypothetical interaction effect.

### Information score

PID information measurement has been proved to be applicable to gene regulatory network inference(25,26); a simple generalized model example was discussed in the work of Timme *et al*.(24). We quoted this example to illustrate the information distribution of PID in different network models. As shown in Figure 1D, let *X*_*1*_, *X*_*2*_ and *Y* be the first source variable, the second source variable and the target variable, respectively; the connection probabilities between the nodes were determined. As a result, we found that in these two network models of the Ex.1 and Ex.2, the redundant and synergistic contribution provided about the target variable by two source variables is high, but the unique information is low or even zero, even though the fact that the two source variables are directly influencing the target variable. In contrast, the other two network structures of Ex.3 and Ex.4 illustrated that most of the information provided about the target variable by the source variables is unique information; the redundant and synergistic contribution is small or even zero. Our three-variable network structure, i.e., miRNA-mRNA-miRNA, is of the same type as Ex.1. We tested it using TCGA breast cancer data (Figure 1D [bottom]). Let *X*_*1*_, *X*_*2*_ and *Y* be the expression of miR-34a-5p, miR-34b-5p and MYC, respectively. The result showed that the information about *Y* provided by *X*_*1*_ and *X*_*2*_ was synergy (Figure 1D [middle bottom]). We further tested alternate combinations of source and target variables in this network structure to explore the underlying law of different information distribution generated by PID. If *X*1 was replaced by *X*2, the information about *Y* provided by *X*_*1*_ and *X*_*2*_ was all redundancy, and there is no unique and synergistic information (Figure 1D [left bottom]. If *X*1 was a copy of *Y*, the total information about *Y* provided by *X*_*1*_ and *X*_*2*_ is the unique information from *X*_1_ (Figure 1D [right bottom]. Thus, we envisage employing PID information measure to interpret both the range and possible underlying mechanism of interaction effects among miRNAs. In our case, we focused on the statistics of how two source variables *R*, i.e., miRNA1 and miRNA2, separately or jointly specify their common target variable *Y*, i.e., mRNA. According to the previous analysis of Figure 1D, the structure of our network suggests that synergistic and redundant interactions between two miRNAs are not mutually exclusive; the magnitude of unique information from either miRNA1 or miRNA2 is negligible compared to the synergistic and redundant information. The high value of synergistic information implies that two miRNAs together can provide most of the information to the target mRNA, while the high value of redundant information means that either miRNA from source set can specify the target mRNA independently. Indeed, unique information is best regarded as a degenerate form of redundancy or synergy in this case(17), so synergy and redundancy alone constitute the basic building blocks of multivariate information. Thus, we derive an information score called ‘synergy-redundancy’ score for each miRNA-mRNA-miRNA triplet to define the direction of interaction effect between two miRNAs, where a positive value indicates a possible synergistic interaction effect and a negative value indicates a potential competitive interaction effect.

## AVAILABILITY

The sycomore is an open-source R program with documented functions and walk-through examples described in the vignette. Main functions were implemented by referring to an R/Bioconductor package we developed earlier called Informeasure. The sycomore is available in the GitHub repository (https://github.com/chupan1218/sycomore)

## RESULTS

### Neighbouring and overlapping binding sites at the sequence level

We analysed the seed matching site data of miRNA targeting mRNA publicly available from TargetScan. TargetScan provides conserved 8mer,7mer and 6mer binding sites that match the seed region of each miRNA(2). Through data cleaning, the database includes 149314 miRNA-mRNA interactions, involving a total of 213 miRNAs and 10458 mRNAs. On average one miRNA is predicted to target 701 mRNAs, and one mRNA is predicted to targeted by 14 miRNAs. We partitioned miRNA pairs into two groups: neighbouring and overlapping, by referring the miRNA target location of seed matching site. The two heatmaps of Figure 2A and 2B respectively illustrate the number of target mRNAs on which miRNA pairs have neighbouring and overlapping binding sites. We noticed that potential neighbours among miRNA target sites are more prevalent than overlaps. There are 3716407 neighbours and 399812 overlaps, of which 63352 pairs are found to have both overlapping and neighbouring loci (Figure 2C). Specific to the number of binding sites per miRNA-mRNA-miRNA triplet, we found that the number of sites is concentrated in the range of less than 3 either in the neighbouring or in the overlapping group. Figure 2D(left) shows that most triplets in neighbouring group have <3 binding sites; only a few have >5 binding sites. Alternatively, Figure 2D(right) shows that most triplets in overlapping group have <=2 binding sites; only a few have >3 binding sites. After the Lasso operation, each cancer-specific miRNA target site landscape also follows the rule that most miRNA pairs are enriched with a small number of binding sites in both neighbouring and overlapping group (Figure 2E). All these statistical evidences show that there indeed exists prevalent neighbouring and overlapping between miRNA target sites.

**Fig. 2.**
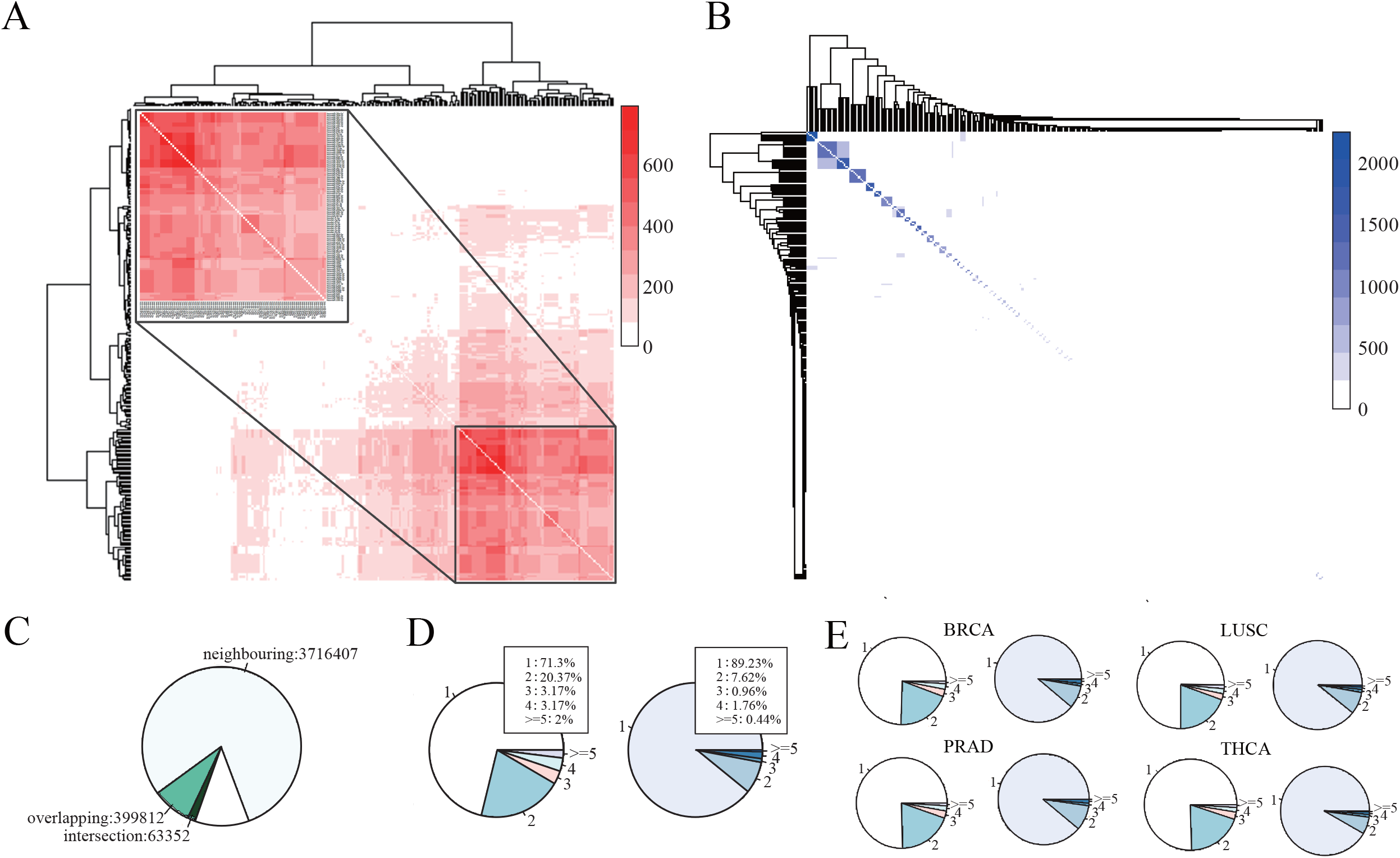
Summary of miRNA target binding sites. **A** miRNA pairs that have neighbouring binding sites. The neighbouring pairs in the upper left corner are magnified in the inset. **B** miRNA pairs that have overlapping binding sites. Colour of each cell in both **A** and **B** represents the number of mRNAs on which a miRNA pair has neighbouring or overlapping binding sites. **C** Number of binding sites includes neighbouring, overlapping and the intersection. **D** Statistics on the number of neighbouring (left) and overlapping (right) binding sites in triplets. **E** Statistics on the number of neighbouring (left) and overlapping (right) binding sites in triplets for each cancer.

### Interaction effects at the expression level

To demonstrate our informatics method, we extensively exploited four cancer RNA-Seq expression profile data downloaded from the TCGA portal: breast invasive carcinoma (BRCA), Lung squamous cell carcinoma (LUSC), prostate adenocarcinoma (PRAD) and Thyroid carcinoma (THCA). We applied PID measurement to each candidate triplet to evaluate an information score list: synergy, redundancy and two unique information. By computing the magnitude of the ‘synergy-redundancy’ score (i.e., information score), we determined which force prevails in the contest of synergy and redundancy. We also estimated miRNA pair co-expression correlation and conducted gene differential expression analysis to jointly explore possible causes for such implicit mechanisms between miRNAs.

### Triplets with negative ‘synergy-redundancy’ scores

We first examined those cases where the ‘synergy-redundancy’ scores were negative, as the number of triplets that met this filter condition was not large. All result lists can be approached in the supplement material. Several intriguing results were found in this examination. Figure 3A illustrates the detailed information distribution of 10 triplets with negative ‘synergy-redundancy’ scores cross four cancer data. On the whole, most of the information provided about the target mRNA by two miRNAs is synergy and redundancy; the two unique information contributions are small or even zero. While the negative ‘synergy-redundancy’ scores imply that the potential synergy effects are more powerful than possible competitive effects, the actual quantitative differences are not large. Of these triplets, two are from BRCA, one is from LUSC, three are from PRAD and the remaining four are from THCA. Among them, only the triplet of miR-141-3p, miR-200c-3p and TRHDE from BRCA has one neighbouring binding site; the others have one overlapping binding site (Figure 3B). Interestingly, most of these miRNA pairs are either from the same miRNA family or are reported to belong to the same miRNA cluster. Typical examples are miR-141-3p, miR-200a-3p, miR-200c-3p and miR-429 from miR-200 family(27), miR-221-3p and miR-222-3p from miR-221/222 cluster(28), miR-17-5p and miR-20a-5p from miR-17/92 cluster(29) (Figure 3B). Others such as miR-106b-5p and miR-93-5p from miR-106b-25 cluster(30), and miR-25-3p and miR-92a-3p from miR-92a family(31) (Figure 3B). Note that the miR-92a family is a group of highly conserved miRNAs including miR-25, miR-92a, miR-363 arise from three homologous miRNA clusters, namely, miR-106-25, miR-106a-363 and miR-17-92(31). Moreover, all miRNA pairs from triplets with negative ‘synergy-redundancy’ scores show a high degree of co-expression correlation (Pearson cc = 0.885 on average, Figure 3C).

**Fig. 3.**
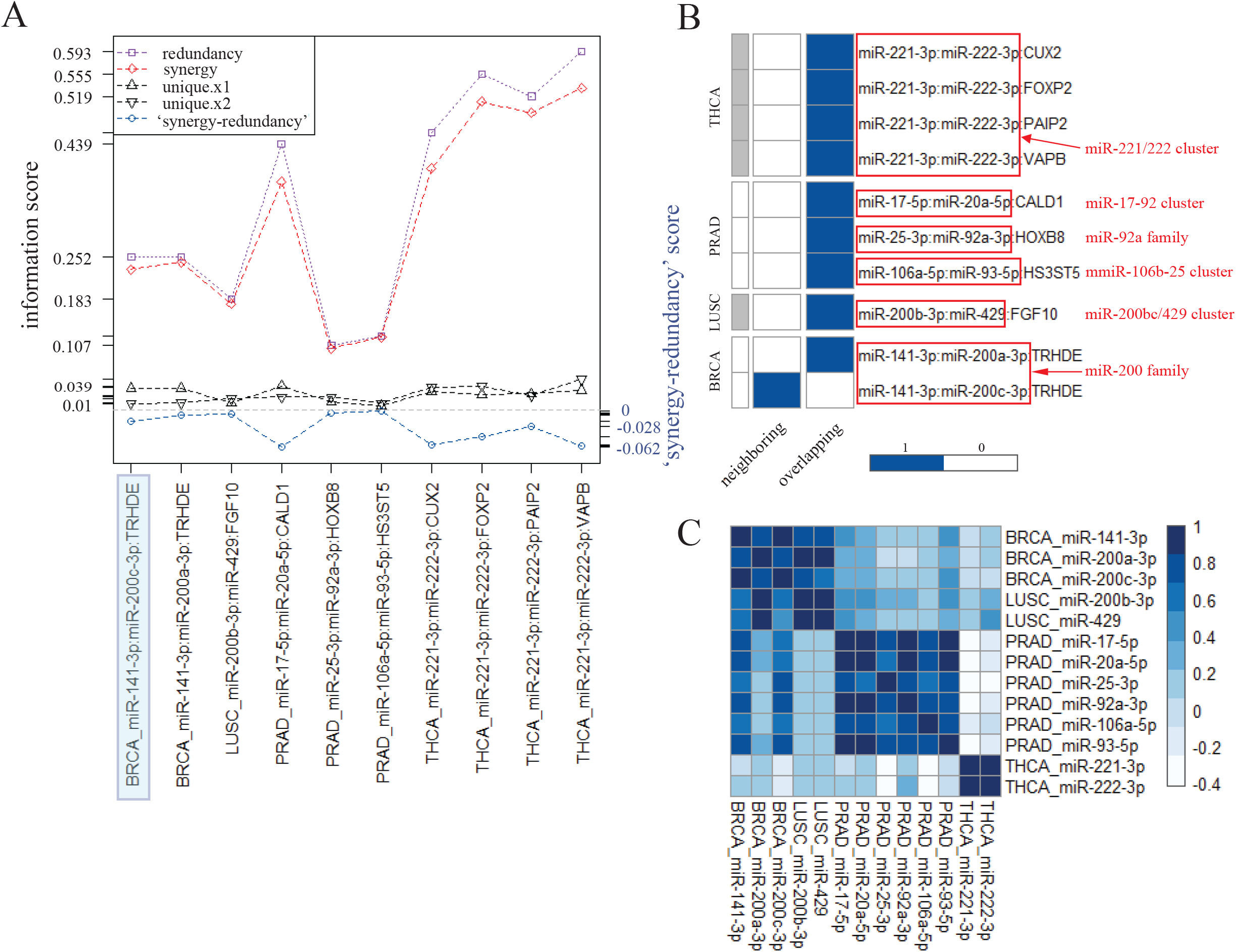
Analysis of these triplets with negative ‘synergy-redundancy’ scores. **A** Information distribution. The shaded area marks that the triplet only has one neighbouring miRNA target site; the others have one overlapping miRNA target site. **B** Number of miRNA target sites per triplet. Tag the miRNA family/cluster to which the miRNA pair belongs. **C** Co-expression correlation of miRNA pairs across four cancer data.

### Triplets with positive ‘synergy-redundancy’ scores

We then examined these cases with positive ‘synergy-redundancy’ scores. All result lists also can be approached in the supplement material. Several intriguing results were observed in this examination. Figure 4 illustrates the analysis of these triplets in BRCA and LUSC; the other two cancers (i.e., PRAD and THCA) can be referred to the supplement figure (Figure S1). We limited the triplets only to those whose Pearson cc corresponding to miRNA pairs are greater than 0.5. As a result, the overall information distribution of these triplets shows that the synergy values are much greater than the redundancy values; two unique information contributions are small, or even zero (Figure 4A, B [left top]). We arranged these triplets in descending order according to the magnitude of Pearson cc of the miRNA pair, accompanied the information scores and the fold change values of triplets (Figure 4A, B [right top]). We found that the top-ranked triplets have low ‘synergy-redundancy’ scores, and the corresponding miRNA pairs show consistent expression patterns between tumour and normal samples, but are different from the targets. While with the gradual decrease of Pearson ccs, the corresponding ‘synergy-redundancy’ scores gradually increased. We verified that Pearson ccs are negatively associated with the ‘synergy-redundancy’ scores (Figure 4A, B [left bottom]). We also checked the miRNA target sites for these triplets (Figure 4A, B [middle bottom]). The binding site statistics tables display that all these triplets have at least one neighbouring binding site; a few have overlapping binding site(s). We further examined the expression correlation of miRNA pairs by adapting the k-mean clustering method. Intriguingly, miRNA pairs from the top 30 triplets are usually partitioned into the same cluster, showing a high co-expression correlation between two miRNAs (Figure 4A, B [right bottom]). Similar to the cases with negative ‘synergy-redundancy’ scores, these miRNA pairs are also from the same miRNA family or are reported to belong to the same miRNA cluster. For example, miR-200a-3p, miR-200b-3p and miR-429 are from miR-200a/200b/429 cluster(27), miR-17-5p, miR-19a-3p and miR-20a-5p from miR-17-92 cluster(29), miR-181a-5p and miR-181b-5p from miR-181 family(32), miR-224-5p and miR-452-5p from miR-224/452 cluster, miR-96-5p and miR-182-5p from miR-183/182/96 cluster(33) (Figure 4A, B [right bottom]).

**Fig. 4.**
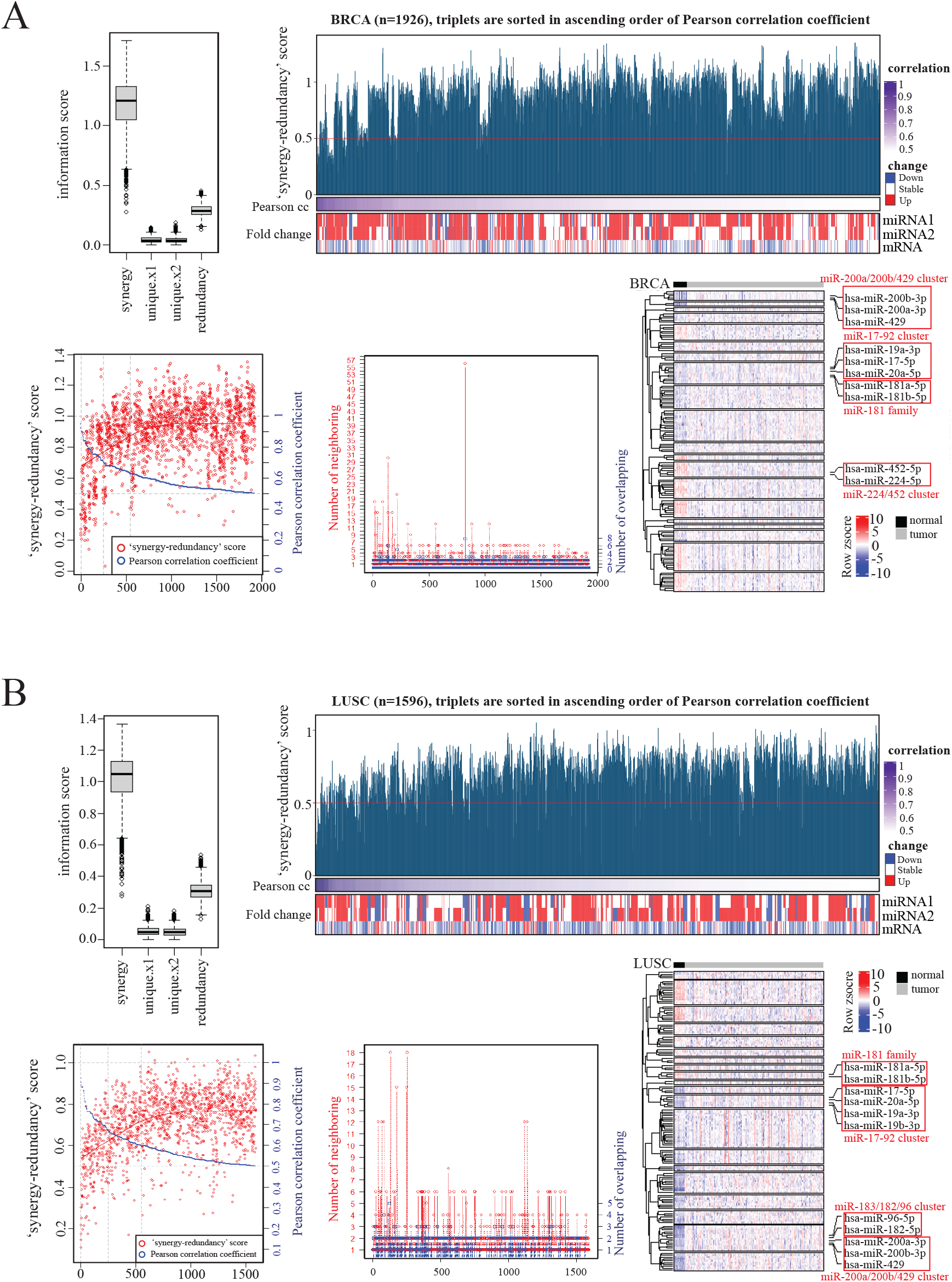
Analysis of these triplets with positive ‘synergy-redundancy’ scores in breast cancer, and lung cancer. **A** (Top left) Information distribution for BRCA data. (Top right) An overview of the association between ‘synergy-redundancy’ scores and co-expressions and expression patterns in BRCA. The triplets are arranged in descending order according to the magnitude of Pearson correlation coefficient of the miRNA pair, accompanied the information scores and the fold change values of triplets. (Bottom left) Correlation of ‘synergy-redundancy’ scores and the Pearson correlation coefficients of the miRNA pairs. (Bottom middle) Number of miRNA target sites per triplet. (Bottom right) Heatmap of miRNAs at BRCA expression level. K-means algorithm was used for partitioning the miRNA clusters. **B** Similar to **A**, examined in LUSC data.

## DISCUSSION

In this article, we proposed a novel computational method called sycomore to quantify the synergy and competition between two miRNAs when targeting on the same mRNA. It uses the information transmission of source variables to the common target variable to quantify the interaction effects between two source variables. We interrogated TargetScan and TCGA data to preliminarily approach the possible mechanism of such implicit interaction effects. Through the sequence data analysis, we found that there indeed exists prevalent neighbouring and overlapping between miRNA target sites, which provides the possibility for miRNAs trigging synergistic or competitive interaction effects. Through the expression data analysis, we noticed that in both cases those triplets with low absolute magnitude of ‘synergy-redundancy’ scores usually have high co-expressions for the corresponding miRNA pairs, where high co-expression usually means high functional similarity between two miRNAs. One possible explanation is that the information passed from highly similar source variables (i.e., miRNA pairs) to common target variable (i.e., mRNA) includes a large amount of synergistic information and huge redundant information as well. Redundant information is defined as the minimum information about the target variable provided by any source, averaged over all possible outcomes(17), so the large amount of redundant information captured means a high information overlap between source variables, which in return implies a high degree of similarity between source variables. This is why the Pearson ccs are negatively associated with the ‘synergy-redundancy’ scores, because as Pearson ccs decreases, the similarity of its corresponding miRNA pairs decreases, thereby reducing the ability to generate redundant information. In fact, miRNA family/cluster analysis also confirmed that miRNA pairs with high functional similarity are likely to derive synergistic and redundant information.

According to the above analysis, we concluded three aspects are important in determining the interaction effects between two miRNAs when targeting on the same mRNA transcript. On is the physical miRNA target sites; another is the co-expression correlation between two miRNAs; the third is the information score (i.e., ‘synergy-redundancy’ score) of the triplet. Sequence is the basis, but not the determining factor. There are a number of triplets with neighbouring binding sites, but few exhibit synergistic interaction effects; similarly, there are number of triplets with overlapping binding sites, but only a few exhibits competitive interaction effects. High co-expression correlation is an essential precondition, but also not the determining aspect. There are several miRNA pairs with high co-expression correlation but show different interaction effects such as miR-200b-3p and miR-429 in LUSC. The positive/negative information score is necessary, but it is still impossible to distinguish between synergy and competition alone. There is a triplet from BRCA with negative ‘synergy-redundancy’ score but presents one neighbouring binding site. However, we find these three aspects together can determine whether the two miRNAs exhibit synergistic or competitive interaction effects when targeting the same mRNA. We thus concluded that the miRNA pair with neighbouring binding site(s) on the mRNA is likely to trigger synergistic events, while the miRNA pair with overlapping binding site(s) on the mRNA is likely to cause competitive events, provided that the pair of miRNAs has a high functional similarity and the corresponding triplet presents a positive/negative ‘synergy-redundancy’ score. These findings may help to enrich the miRNA-mRNA regulation landscape.

## SUPPLEMENTARY DATA

**Supplementary Data** are available at NAR online.

## ACKNOWLEDGEMENTS

We would like to thank Zhaolei Zhang, Zhonghao Wang, Tingting Hu and Mingyi Zhao for helpful discussion.

