## Supplementary figures and images for "Explore synergistic and competitive miRNA regulation mechanisms in the miRNA-mRNA regulatory network from the information decomposition perspective"

### Figure S1.pdf

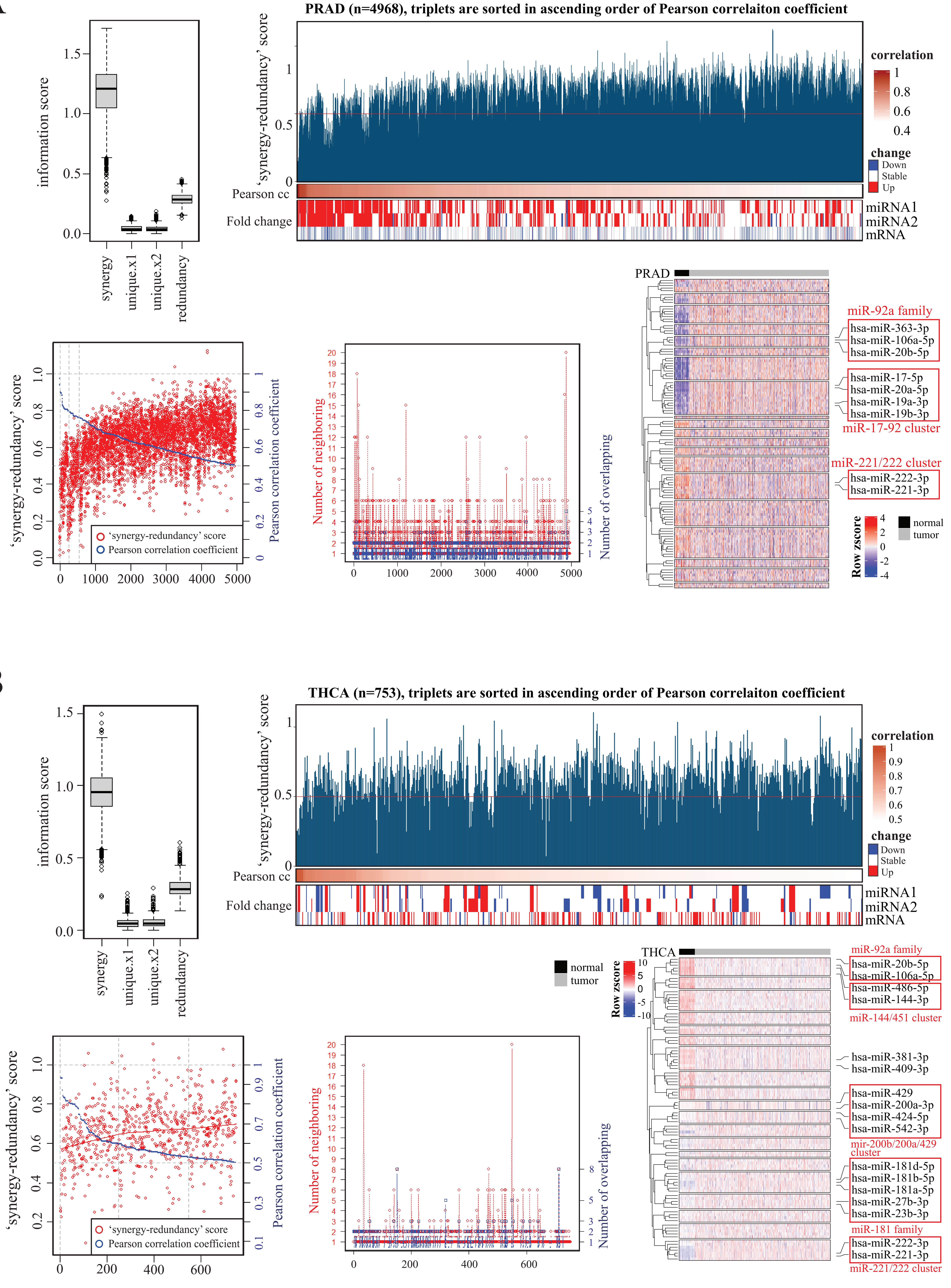
